# Molecularly Targeted Magnetic Resonance Imaging and Spectroscopy

**DOI:** 10.1101/2020.04.05.026252

**Authors:** Jia-Xiang Xin, Yi Li, Hui-Xia Liu, Jianqi Li, Guang Yang, Huojun Zhang, Jiachen Wang, Rui Tong, Da-Xiu Wei, Ye-Feng Yao

## Abstract

Magnetic resonance imaging (MRI) and magnetic resonance spectroscopy (MRS) have tremendous utility in many fields, such as clinical diagnosis, medical research and brain science. MRI provides high resolution anatomic images of tissues/organs, and MRS provides functional molecular information related to specific regions of tissues/organs. However, it is often difficult for conventional MRI/MRS to selectively image/probe a specific metabolite molecule other than water and fat. This greatly limits study of the molecular mechanisms underlying metabolism and disease. Herein, we report a novel method for obtaining an exact molecularly targeted MRI and MRS. This method uses the nuclear spin singlet state to select the signals from a specific molecule. Several endogenous molecules in living organism such as N-acetylaspartate and dopamine have been exemplarily imaged and probed as the targeted molecules in the MRI and MRS experiments, demonstrating the unique molecular selectivity of the developed method.

**Endogenous-molecule-targeted MRI and MRS can be achieved by using the new pulse sequences:** 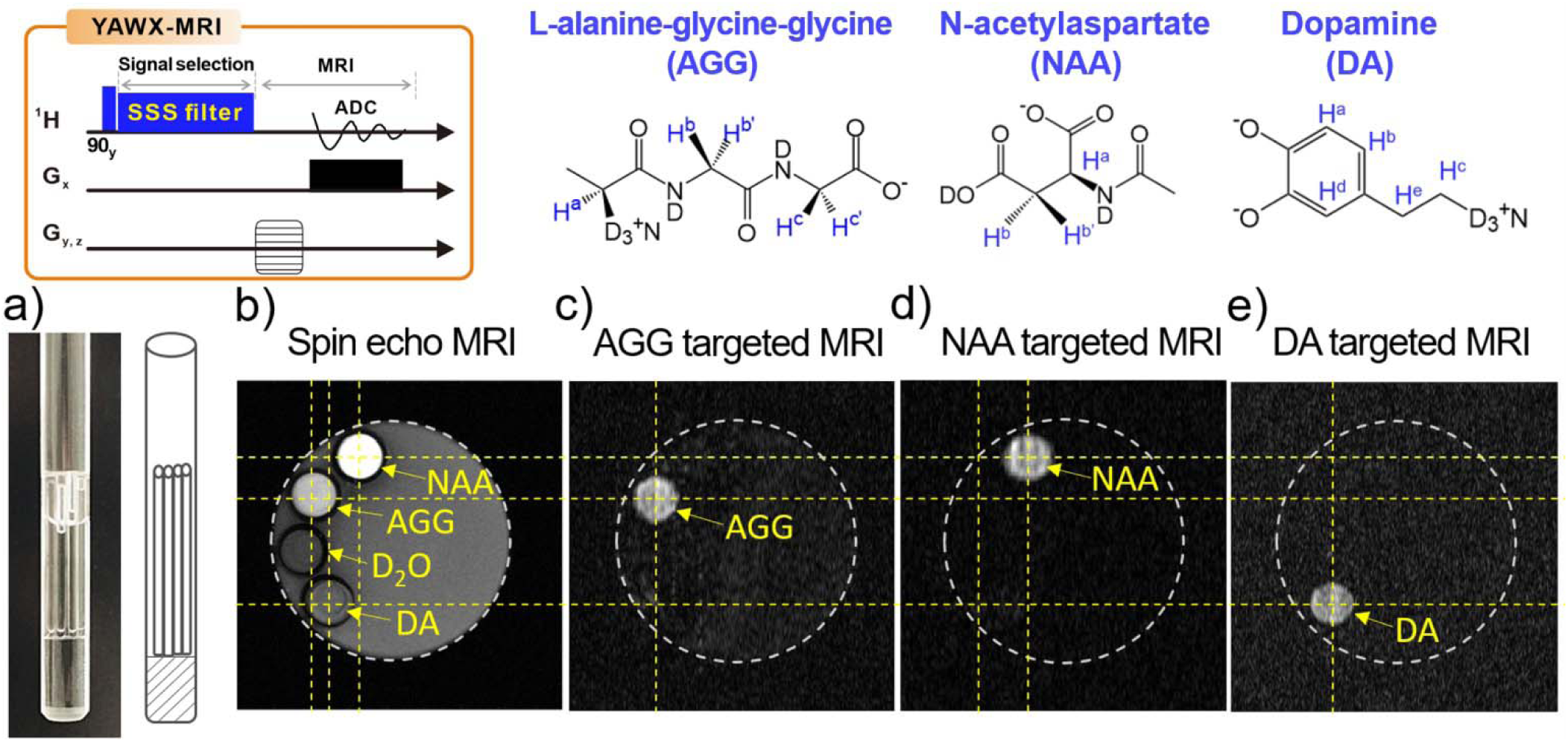

## INTRODUCTION

Magnetic resonance imaging (MRI) is widely used in clinic diagnosis, biomedical research, and brain science^1-4^. Most MRI techniques map and measure ^1^H spins from water and fat^5^. Although MRI can detail *in vivo* anatomy with very high spatial resolution, conventional MRI techniques often lack “chemical resolution”: They cannot selectively image/probe a specific metabolite other than water^1-3^. Magnetic resonance spectroscopy (MRS) is complementary to MRI and can directly probe different biosubstances^3, 6-7^. A routine MRS procedure includes the acquisition of MR images and the corresponding MR spectra. The MR images offer anatomic landmarks that correspond to the MRS data, while the MRS data provide the molecular information of MR image. This combination of anatomy and molecular information facilitates accurate diagnosis, recognition, and therapy^8-12^. However, the Achilles’ heel of MRS is the signal overlap problem due to the complexity of bio-substances, which results in great difficulty to analyze and quantify MRS signals^6-7, 13^. Selective imaging/detection of endogenous molecules is in dire need of various MRS practical applications^14-18^.

Many efforts in MRI and MRS have been made along this direction. In MRI, exogenetic biomarker or targeted agents have been developed to achieve an imaging of a specific molecule^17-20^. By exploiting the chemical exchange process between hydroxyl groups of water and the active hydrogens of the biosubstances, Balaban et al. developed the chemical exchange dependent saturation transfer (CEST) method to achieve an indirect detection of the biosubstances such as amino acid and sugars^21^. In MRS applications, various spectral editing techniques, such as MEGA editing by Mescher and Garwood et al.^22^, double quantum filtered (MQF) MRS^23-25^, have been developed to selectively probe the signals from the specific biomolecules. To probe the spatial distributions of the metabolites in organs, Brown et al. developed magnetic resonance spectroscopic imaging (MRSI)^26^. Although these techniques can selectively probe molecules, they fail to accurately select a specific molecule for true molecularly targeted imaging/detection.

In this work, we will demonstrate that by exploiting the nuclear Spin Singlet State (SSS) we are able to achieve true molecularly targeted MRI/MRS. The SSS is defined as a nuclear spin state with a relaxation time much longer than the longitudinal relaxation time T_1_. Thus, it is often called the long-lived nuclear spin singlet state^27-28^. The long-lived nature of SSS offers the opportunities to probe extremely slow molecular motion^29-32^ and chemical exchange^33^, to increase the contrast of magnetic resonance spectroscopy and imaging^31, 34-35^, and to increase the coherent time in quantum computation^36^ and so on^37-39^. Especially, it has been presented that SSS can be used as a filtering method to select signals from a specific molecule, and then some endogenous molecules in living organism can be exemplarily imaged or probed as the targeted molecules^40-47^. Very recently, Glöggler’s group demonstrated the application of SSS filtering method in the Alzheimer’s disease-related b-amyloid 40 peptide and in metabolites in brain matters^48^. However, another intriguing virtue of SSS, which importance has not gotten enough attention, is its instinctive immunity to the gradient field during spin coherence evolution^49-50^. By exploiting this property, the evolution period of SSS can be used to select the signals from SSS while suppressing all of the other signals by applying a gradient field (see the supplementary text in Supporting Information for more details). Moreover, the preparation of SSS of a target molecule, which requires the unique combination of the chemical shift difference and J-coupling^28, 51-52^, introduces the salient molecule targeting feature to this technique. Combination of the molecule targeting feature and the immunity to the gradient during evolution makes the SSS-based techniques possess the perfect molecular level selectivity.

## RESULTS AND DISCUSSION

Figure 1 shows three pulse sequences designed for molecularly targeted signal selection in nuclear magnetic resonance (NMR), MRI, and MRS applications. Figure 1a shows the **YA**o**W**ei**X**in-NMR pulse sequence (YAWX-NMR) that can select signals from a specific molecule in NMR applications. The core part of this pulse sequence is the SSS filter that consists of three blocks in sequence: The first spin lock sequence block is used to prepare the singlet state^52^. The second block consists of two gradient pulses and a decoupling pulse between them to select the singlet signals while suppressing all other signals. The third block is a spin lock sequence block that transforms the singlet signals into observable signals. The details of the spin evolution in YAWX-NMR can be found in the supplementary text in Supporting Information.

**Figure 1:**
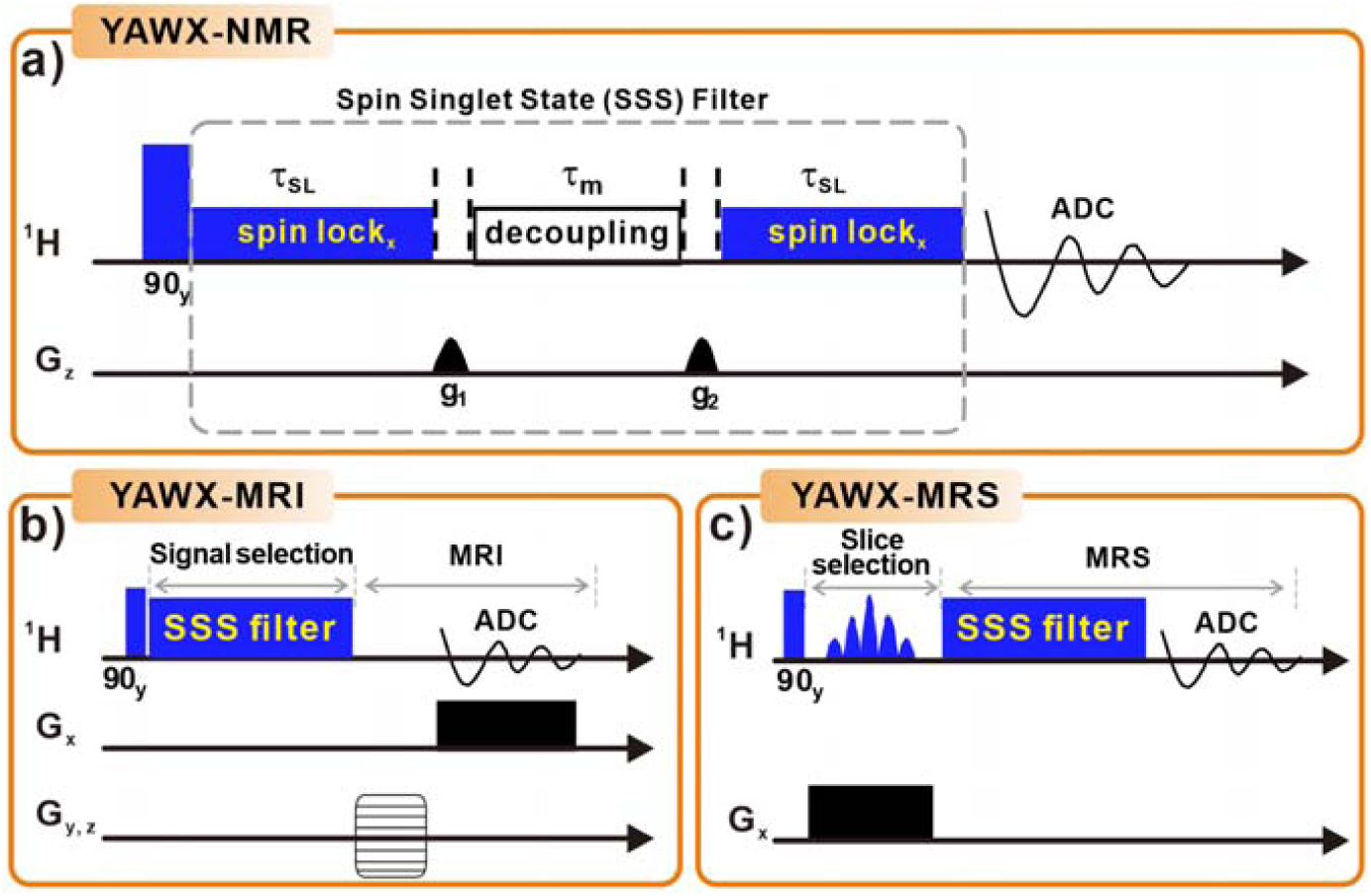
The scheme of the pulse sequence designed for: a) the molecular targeted NMR, **YA**o**W**ei**X**in-NMR, Abbr. YAWX-NMR, b) 2D-MRI via YAWX-MRI, and c) MRS via YAWX-MRS. In YAWX-NMR, the blue filled rectangles represent the 90° pulses and the spin lock block, respectively. The empty rectangle represents the decoupling block. The pulsed-field gradients, g_1_and g_2_, are used for coherence dephasing. In YAWX-MRI, the blue filled rectangles represent the 90° pulses and the SSS filter, respectively. The pulsed-field gradients are used for imaging. In YAWX-MRS, the blue filled rectangles represent the 90° pulses and the SSS filter, respectively. The blue shaped pulse and the slice-selection gradient (the black filled rectangle) are used for slice selection. The detailed experimental parameters can be found in Supporting Information.

The SSS filter can be used in MRI and MRS sequences to achieve molecular selectivity in different applications. Figure 1b shows the YAWX-MRI pulse sequence for molecularly targeted MRI. This pulse sequence consists of the SSS filter block to achieve signal selection of the targeted molecule as well as 3D imaging of the sequence block. Figure 1c shows the YAWX-MRS pulse sequence that is designed for molecularly targeted localized MRS. In this pulse sequence, a gradient spin echo is used for spatial slice selection, and the SSS filter block selects the signal from a target molecule. More details of these pulse sequences can be found in Supporting Information. Note that the key idea underlying these three pulse sequences is the use of a SSS for signal selection. We note that many pulse sequences have been reported for the preparation of SSS in the literature^28-31, 36-39, 49^, and these pulse sequences can theoretically all be used as SSS filters.

The molecular selectivity of SSS filters originates from the strict preparation condition of the SSS. The core parameters of the SSS filter (e.g., the duration time and the power level of the spin lock) are calculated from the properties of the spin system (i.e., the chemical shifts and J coupling). The one-to-one relationship between the core parameters of the SSS filter and the properties of the spin system is critical to the molecular selectivity of the SSS filter. In principle, any molecule with a spin system suitable for SSS preparation can be used as a target molecule in the SSS filter. Such spin systems can be two^49, 53^ or multiple coupled homonuclear spins^54-57^, isolated or non-isolated from other nuclei spins^58-59^. Generally, it is easier to prepare a high-transfer-efficiency long-lived state in a two-spin system than in a multi-spin system^49, 53^. The transfer efficiency of long-lived singlet state prepared from an isolated spin system is higher than that from a non-isolated spin system. Note that the long-lived singlet state of a non-isolated spin system can be prepared and used in the SSS filter, although in such cases the relaxation attenuation of the long-lived state would be accelerated. This greatly extends the choice of targeted molecules. Some important biochemicals, such as N-acetylaspartate (NAA) and dopamine (DA) can be used as the target molecule in the SSS filter (see Figure S2 in Supporting Information). This provides a very promising application of the SSS filter to selectively probe biochemicals in living organisms *in vivo*. In the following discussion, three target molecules - the tripeptide L-alanine-glycine-glycine (AGG^51^), NAA, and DA - will be used as examples to demonstrate the molecular selectivity of YAWX-NMR/MRI/MRS.

Figure 2a shows the ^1^H spectrum of the AGG aqueous solution acquired using a single pulse excitation. Figure 2b shows the ^1^H spectrum of the AGG aqueous solution acquired with YAWX-NMR. The singlet state of the two-spin system (H^c^and H^c’^) was prepared herein. The preparation of the SSS of H^c^and H^c’^requires a strict match between the parameters in the pulse sequence and the properties of the spin system (i.e., the chemical shift and J coupling of the spin system). Therefore, only the signals from H^c^and H^c’^ can pass through the SSS filter, whereas all other signals (including the water signals) were filtered out or significantly suppressed. To further demonstrate the excellent signal selectivity, we performed YAWX-NMR on the aqueous solution of the mixture of AGG and insulin. The ^1^H spectra acquired using a single pulse excitation and YAWX-NMR are shown in Figure 2c and 2d, respectively. Similar to the case in Figure 2a and 2b, only the signals from H^c^and H^c^’ of AGG and the significantly suppressed water signal remain in the spectrum, although insulin molecules give many resonance signals in the spectrum.

**Figure 2:**
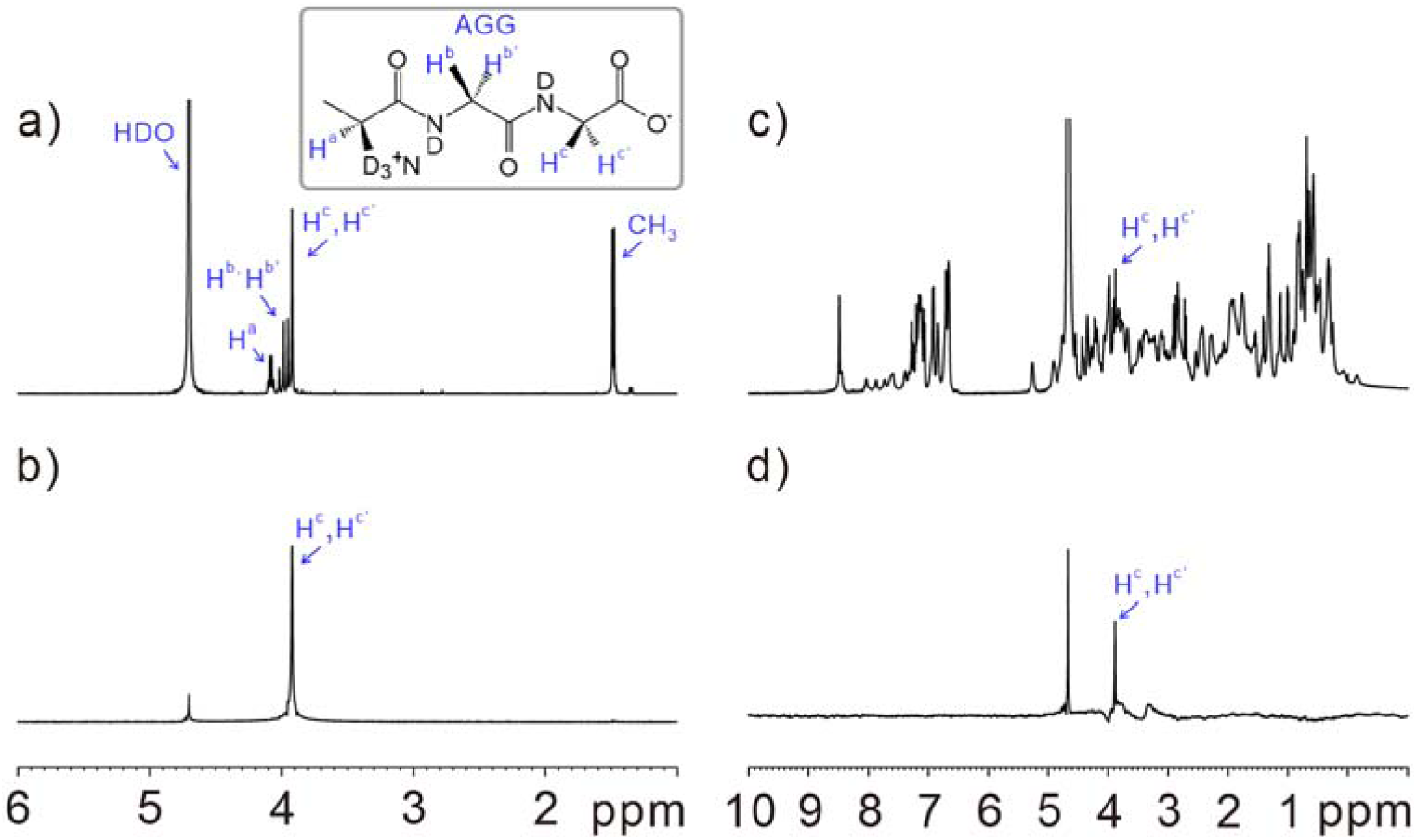
The ^1^H NMR spectra of the aqueous solution of a tripeptide (L-alanine-glycine-glycine, AGG^51^) was acquired by using: a) the single pulse excitation sequence; and b) the YAWX-NMR pulse sequence to select the signals of H^c^and H^c^’ of AGG. The ^1^H NMR spectra of the aqueous solution of the mixture of insulin and AGG, acquired by using: c) the single pulse excitation sequence; and d) the YAWX-NMR pulse sequence to select the signals of H^c^and H^d^’ of AGG. The experimental temperature is room temperature. The recycle delay in the experiments was set to 10 s to ensure the full relaxation of the signals. More experimental details can be found in Supporting Information.

The SSS filter was also introduced into the MRI sequence to achieve molecularly-targeted MRI. To demonstrate this concept, we designed a special phantom based on a 5 mm NMR glass tube containing water (40% D_2_O and 60% H_2_O). The concentration of D_2_O was adjusted to optimize the signal intensity in the imaging. Four capillaries (1 mm diameter, ∼0.1 mm wall thickness) containing D_2_O (60% D_2_O and 40% H_2_O) as well as the NAA, AGG, and DA aqueous solution were carefully placed in the 5 mm tube. The photo and schematic illustration of this phantom are shown in Figure 3a. Figure 3b shows a routine ^1^H MRI image of the sample acquired using the spin echo imaging sequence. The image shows one large grey disk containing four small disks. Each small disk is surrounded by a black circle. The big grey disk is from the water inside the 5 mm glass tube, and the black, gray, and white disks are from the capillaries containing the HDO water, AGG, NAA, and DA aqueous solutions, respectively. The different signal intensity can be attributed to the different ^1^H concentrations in different solutions. Each small disk has a black ring showing the wall of the capillary tubes. We used YAWX-MRI to obtain molecularly targeted images of the sample. Figures 3c, d, and e show the molecularly targeted ^1^H MRI image of AGG, NAA and DA, respectively, acquired using the pulse sequence in Figure 1b. The selected signals for imaging are H^c^and H^c^’ of AGG, H^b^and H^b^’ of NAA and H^d^of AGG, respectively. The singlet preparation was optimized based on the simulation of the singlet transfer efficiency against the spinlock time and power in Figure S3-5 in Supporting Information. More experimental details are referred to the method part in Supporting Information. The images in Figure 3c, d, and e clearly demonstrate that each capillary containing the corresponding targeted molecule can be selectively imaged indicating the excellent molecular selectivity of the method.

**Figure 3:**
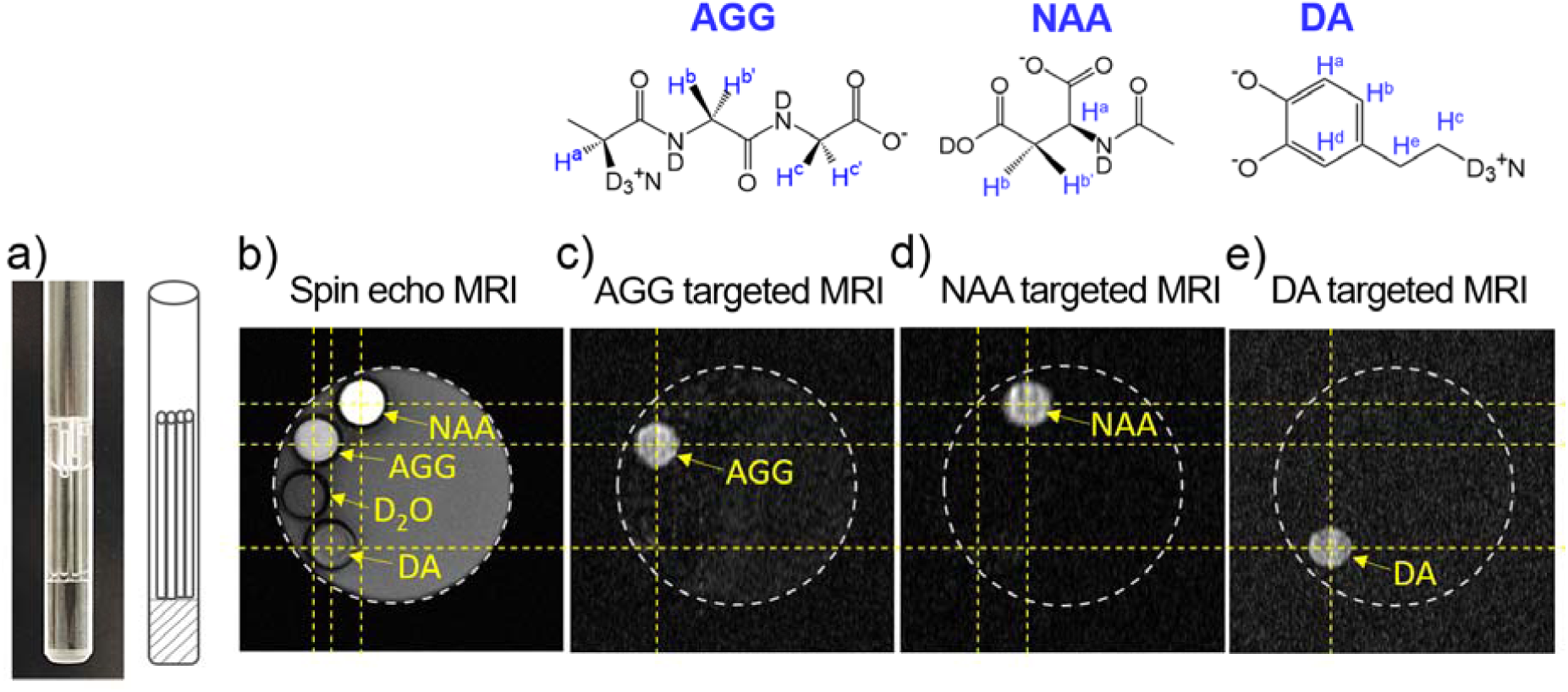
a) A photo of the phantom. This sample is a 5-mm NMR glass tube containing water (40% D_2_O and 60% H_2_O). Four capillary tubes (1 mm diameter) containing D_2_O and the NAA, AGG, and DA aqueous solution respectively, were carefully set into the 5 mm tube. b) The ^1^H MRI image of the sample, acquired by using a standard ^1^H spin echo imaging sequence (T _E_ = 15.5 ms, T _R_ = 10 s, field of view = 5.6 × 5.6 mm^2^); c) TheAGG targeted MRI image of the sample, where CH_2_ signals (H^c^and H^c^’, see Figure 2) of AGG were selected for imaging. d) The NAA-targeted MRI image of the sample where CH signals (H^b^and H^b^’, see Figure 2) of NAA were selected for imaging. e) The DA targeted MRI image of the sample where the CH signal (H^d^, see Figure 2) of DA was selected for imaging. The molecular MRI images of the sample were acquired using the YAWX-MRI pulse sequence in Figure 1b. The experimental temperature is room temperature. The recycle delay in the experiments was set to 10 s to ensure full relaxation of the signals. More experimental details can be found in Supporting Information.

The combination of the SSS filter and MRS can monitor the targeted molecules in specific spatial regions of the sample. Figure 4 shows the molecularly targeted MRS spectra acquired using the YAWX-MRS sequence. The sample was similarly a 5 mm NMR tube with the four capillary tubes containing the HDO, NAA, AGG, and DA aqueous solutions. Figure 4a is the MRI image of the sample. The yellow square indicates the region selected for MRS. Figure 4b is the routine ^1^H MRS spectrum of the selected region in the sample. The signals from the HDO, AGG, NAA, and DA solutions can be observed in the spectrum. The poor resolution and linear shape of the signals is due to the high inhomogeneity of the field due to the disturbance of the inserted capillary tubes.

**Figure 4:**
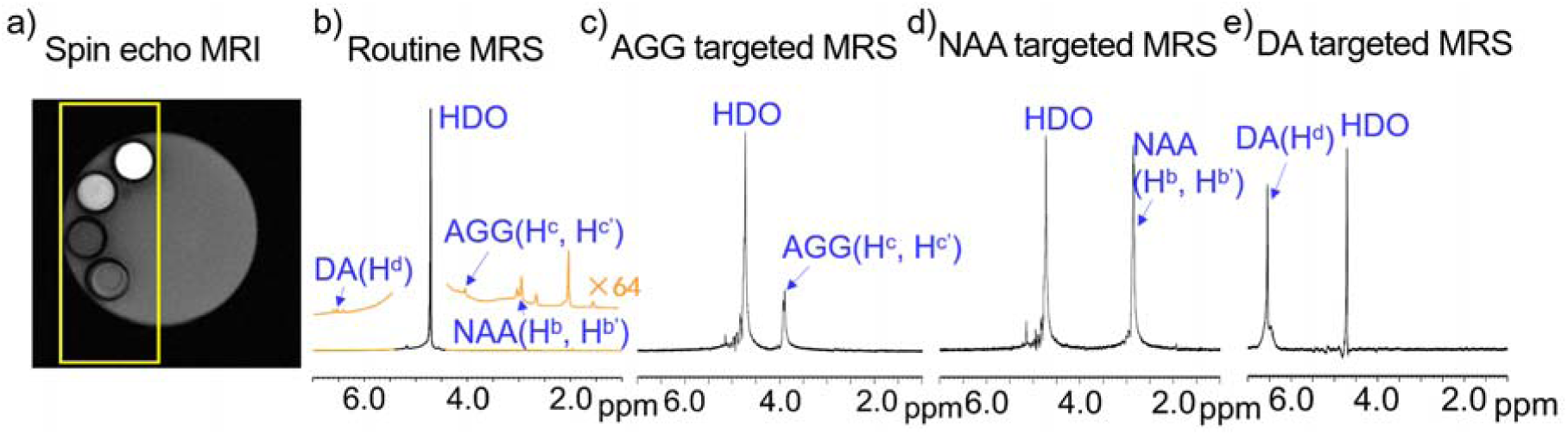
a) The ^1^H MRI image of the sample, acquired by using a standard ^1^H spin echo imaging sequence (T _E_ = 15.5 ms, T _R_ = 10 s, field of view = 5.6 × 5.6 mm^2^). The yellow dash lines indicate the region selected for MRS. b) The routine ^1^H MRS of the selected region in the sample; c) Themolecularly targeted MRS spectrum of the sample. The signals of H^c^and H^c^’ of AGG were detected selectively; d) The molecular targeted MRS spectrum of the sample. The signals of H^b^and H^b^’ of NAA was selectively probed. e) TheDA-targeted MRS spectrum of the sample. The signals of H^d^of DA were selectively probed. The signal selection can have a strong suppression of the water signal. The residual signal of water is about 0.1–0.5% of the original one. The molecularly targeted MRS spectra of the sample were acquired using the YAWX-MRS sequence. The yellow square indicates the region selected for MRS. The experiment was conducted at room temperature. The recycle delay in the experiments was set to 10 s to ensure full relaxation of the signals. More experimental details can be found in Supporting Information.

Figure 4c shows the AGG-targeted ^1^H MRS spectrum. The signals of H^c^and H^c^’ of AGG are selected and remain in the spectrum, whereas the other signals including the water signal are significantly suppressed. Similarly, the signals of H^b^and H^b^’ of NAA and H^d^of DA are dominant in the NAA-targeted and DA-targeted ^1^H MRS spectra in Figures 4d and 4e, respectively. These data indicate the excellent molecular selectivity of the method. Combining the spatial information from the MRI imaging, we are able to use YAWX-MRS to probe the spatial distribution of the target molecules. By using a suitable reference signal, it is also capable to quantify the amount of the target molecule in a specific area, which is the on-going research in our laboratory. Note that compared with the molecularly targeted MRI, the molecularly targeted MRS is a more sensitive and efficient approach to probe the signal of targeted molecule. In Figure S6 in Supporting Information, we show that even for the DA solution having the concentration of 26 μM, the signal, H^d^, of DA still can be easily selectively probed.

## CONCLUSIONS

In this work, we developed a novel method for obtaining an exact molecularly targeted MRI and MRS. This method uses the nuclear spin singlet state to select the signals from a specific molecule. We designed three types of pulse sequences, namely YAWX-NMR, YAWX-MRI and YAWX-MRS, which can be used for the molecular targeted signal selection in the NMR, MRI and MRS applications. To demonstrate the unique molecular targeting feature of the developed method, several important endogenetic molecules in living organism such as NAA and DA have been exemplarily probed and imaged as the targeted molecules in the NMR, MRI and MRS experiments.

We believe that the developed method in this work will have profound influence on cell/molecular biology, brain science, medicine, pharmacology, medical physics, chemistry, physics, MRI/NMR methodology and might inspire many innovative ideas in bio-NMR/MRI. For example, molecular imaging of disease biomarkers using MRI requires targeted MRI contrast agents with high specificity and high relaxivity. While peptides, antibodies, or small ligands can be used to achieve targeting, they must be linked to MRI contrast containing Gd or micron-sized particles of iron oxide (MPIO). Such media not only complicate the synthesis process but also have safety concerns. Our selective imaging/probing approach can directly image a specific endogenetic biomarker, thus eliminating the need for exogenous contrast agents. Even when selective imaging of the biomarker is not possible directly, we can still use external targeting agents without binding them to MRI contrast agents. This method will open up many new possibilities for molecular MRI/MRS, which in turn will facilitate the study of molecular process in living organisms to improve the early diagnosis of diseases and optimize pre-clinical and clinical testing of new medications.

## ASSOCIATED CONTENT

### Supporting Information

Supplementary text, materials, experimental details and the molecularly targeted ^1^H spectra of N-acetylaspartate and dopamine (26.1 mM and 26.1 μM).

## AUTHOR INFORMATION

### Author Contributions

Y. Y. conceived the idea. Y. Y., D. W. and J. X. designed the pulse sequences. J. X., Y. L. and H. L. prepared samples and performed NMR/MRI experiments. J. L., G. Y., H. Z. and J. W. were involved in discussion. All authors co-wrote the manuscript. All authors have read the manuscript and agreed to its contents.

## ACKNOWLEDGMENT

This work was financially supported by National Natural Science Foundation of China (21574043), Ministry of Science and Technology of the People’s Republic of China (2018YFF01012504) and Microscale Magnetic Resonance Platform of ECNU and the Fundamental Research Funds for the Central Universities.

## REFERENCES

1. Blümich, B.; Blümich; Pauly, Essential NMR. Springer: 2019.

2. Johnson, G. A.; Cofer, G. P.; Fubara, B.; Gewalt, S. L.; Hedlund, L. W.; Maronpot, R. R., Magnetic resonance histology for morphologic phenotyping. J. Magn. Reson. Imaging 2002, 16 (4), 423–429.

3. Gallagher, F., An introduction to functional and molecular imaging with MRI. Clin. Radiol. 2010, 65 (7), 557–566.

4. Vandenberg, J. I.; Kuchel, P. W., Nobel Prizes for magnetic resonance imaging and channel proteins. Med. J. Aust. 2003, 179 (11/12), 611–613.

5. Ma, J., Dixon techniques for water and fat imaging. J. Magn. Reson. Imaging 2008, 28 (3), 543–558.

6. Puts, N. A.; Edden, R. A., In vivo magnetic resonance spectroscopy of GABA: a methodological review. Prog. Nucl. Magn. Reson. Spectrosc. 2012, 60, 29.

7. Mullins, P. G.; McGonigle, D. J.; O’Gorman, R. L.; Puts, N. A.; Vidyasagar, R.; Evans, C. J.; Edden, R. A., Current practice in the use of MEGA-PRESS spectroscopy for the detection of GABA. NeuroImage 2014, 86, 43–52.

8. Weissleder, R.; Mahmood, U., Molecular imaging. Radiology 2001, 219 (2), 316–333.

9. Gambhir, S. S., Molecular imaging of cancer with positron emission tomography. Nat. Rev. Cancer 2002, 2 (9), 683–693.

10. Massoud, T. F.; Gambhir, S. S., Molecular imaging in living subjects: seeing fundamental biological processes in a new light. Genes Dev. 2003, 17 (5), 545–580.

11. Ding, F.; Zhan, Y.; Lu, X.; Sun, Y., Recent advances in near-infrared II fluorophores for multifunctional biomedical imaging. Chem. Sci. 2018, 9 (19), 4370–4380.

12. Tempany, C. M.; McNeil, B. J., Advances in biomedical imaging. J. Am. Med. Assoc. 2001, 285 (5), 562–567.

13. Harris, A. D.; Saleh, M. G.; Edden, R. A., Edited 1H magnetic resonance spectroscopy in vivo: Methods and metabolites. Magn. Reson. Med. 2017, 77 (4), 1377–1389.

14. Gallagher, F. A.; Kettunen, M. I.; Day, S. E.; Hu, D.-E., Ardenkjær-Larsen, J. H.; Jensen, P. R.; Karlsson, M.; Golman, K.; Lerche, M. H.; Brindle, K. M., Magnetic resonance imaging of pH in vivo using hyperpolarized 13 C-labelled bicarbonate. Nature 2008, 453 (7197), 940–943.

15. Golman, K.; Lerche, M.; Pehrson, R.; Ardenkjaer-Larsen, J. H., Metabolic imaging by hyperpolarized 13C magnetic resonance imaging for in vivo tumor diagnosis. Cancer Res. 2006, 66 (22), 10855–10860.

16. Jasanoff, A., MRI contrast agents for functional molecular imaging of brain activity. Curr. Opin. Neurobiol. 2007, 17 (5), 593–600.

17. Lee, T.; Cai, L. X.; Lelyveld, V. S.; Hai, A.; Jasanoff, A., Molecular-level functional magnetic resonance imaging of dopaminergic signaling. Science 2014, 344 (6183), 533–535.

18. Viale, A.; Aime, S., Current concepts on hyperpolarized molecules in MRI. Curr. Opin. Chem. Biol. 2010, 14 (1), 90–96.

19. Lee, J.-H., Huh, Y.-M., Jun, Y.-w., Seo, J.-w., Jang, J.-t., Song, H.-T., Kim, S.; Cho, E.-J., Yoon, H.-G., Suh, J.-S.,, Artificially engineered magnetic nanoparticles for ultra-sensitive molecular imaging. Nat. Med. 2007, 13 (1), 95–99.

20. Weissleder, R., Molecular imaging in cancer. Science 2006, 312 (5777), 1168–1171.

21. Ward, K.; Aletras, A.; Balaban, R. S., A new class of contrast agents for MRI based on proton chemical exchange dependent saturation transfer (CEST). J. Magn. Reson. 2000, 143 (1), 79–87.

22. Mescher, M.; Merkle, H.; Kirsch, J.; Garwood, M.; Gruetter, R., Simultaneous in vivo spectral editing and water suppression. NMR Biomed. 1998, 11 (6), 266–272.

23. Choi, I. Y.; Lee, S. P.; Merkle, H.; Shen, J., Single-shot two-echo technique for simultaneous measurement of GABA and creatine in the human brain in vivo. Magn. Reson. Med. 2004, 51 (6), 1115–1121.

24. McLean, M.; Busza, A.; Wald, L.; Simister, R.; Barker, G.; Williams, S., In vivo GABA+ measurement at 1.5 T using a PRESS-localized double quantum filter. Magn. Reson. Med. 2002, 48 (2), 233–241.

25. Shen, J.; Shungu, D. C.; Rothman, D. L., In vivo chemical shift imaging of γ-aminobutyric acid in the human brain. Magn. Reson. Med. 1999, 41 (1), 35–42.

26. Brown, T. R.; Kincaid, B.; Ugurbil, K., NMR chemical shift imaging in three dimensions. Proc. Natl. Acad. Sci. U. S. A. 1982, 79 (11), 3523–3526.

27. Carravetta, M.; Johannessen, O. G.; Levitt, M. H., Beyond the T 1 limit: singlet nuclear spin states in low magnetic fields. PhRvL 2004, 92 (15), 153003.

28. Carravetta, M.; Levitt, M. H., Long-lived nuclear spin states in high-field solution NMR. J. Am. Chem. Soc. 2004, 126 (20), 6228–6229.

29. Cavadini, S.; Dittmer, J.; Antonijevic, S.; Bodenhausen, G., Slow diffusion by singlet state NMR spectroscopy. J. Am. Chem. Soc. 2005, 127 (45), 15744–15748.

30. Ahuja, P.; Sarkar, R.; Vasos, P. R.; Bodenhausen, G., Diffusion coefficients of biomolecules using long-lived spin states. J. Am. Chem. Soc. 2009, 131 (22), 7498–7499.

31. Pileio, G.; Dumez, J.-N., Pop, I.-A., Hill-Cousins, J. T.; Brown, R. C., Real-space imaging of macroscopic diffusion and slow flow by singlet tagging MRI. J. Magn. Reson. 2015, 252, 130–134.

32. Pileio, G.; Ostrowska, S., Accessing the long-time limit in diffusion NMR: The case of singlet assisted diffusive diffraction q-space. J. Magn. Reson. 2017, 285, 1–7.

33. Sarkar, R.; Vasos, P. R.; Bodenhausen, G., Singlet-state exchange NMR spectroscopy for the study of very slow dynamic processes. J. Am. Chem. Soc. 2007, 129 (2), 328–334.

34. Pileio, G.; Bowen, S.; Laustsen, C.; Tayler, M. C.; Hill-Cousins, J. T.; Brown, L. J.; Brown, R. C.; Ardenkjaer-Larsen, J. H.; Levitt, M. H., Recycling and imaging of nuclear singlet hyperpolarization. J. Am. Chem. Soc. 2013, 135 (13), 5084–5088.

35. Devience, S. J.; Walsworth, R. L.; Rosen, M. S., Nuclear spin singlet states as a contrast mechanism for NMR spectroscopy. NMR Biomed 2013, 26 (10), 1204–12.

36. Roy, S. S.; Mahesh, T., Initialization of NMR quantum registers using long-lived singlet states. PhRvA 2010, 82 (5), 052302.

37. Buratto, R.; Mammoli, D.; Chiarparin, E.; Williams, G.; Bodenhausen, G., Exploring weak ligand–protein interactions by long-lived NMR states: improved contrast in fragment-based drug screening. Angew. Chem. Int. Ed. 2014, 53 (42), 11376–11380.

38. Buratto, R.; Mammoli, D.; Canet, E.; Bodenhausen, G., Ligand–Protein Affinity Studies Using Long-Lived States of Fluorine-19 Nuclei. J. Med. Chem. 2016, 59 (5), 1960–1966.

39. Mamone, S.; Glöggler, S., Nuclear spin singlet states as magnetic on/off probes in self-assembling systems. PCCP 2018, 20 (35), 22463–22467.

40. Xin, J.; Liu, H.; Wei, D.; Yao, Y. Chinese Patent: 201811450785.7, 2018.

41. Yao, Y.; Xin, J.; Liu, H.; Wei, D. Patent: PCT/CN2019/121856, 2019.

42. Yao, Y.; Xin, J.; Li, Y.; Wei, D.; Wang J.; Chinese Patent: 201911249153.9, 2019.

43. Yao, Y.; Xin, J.; Li, Y.; Wei, D.; Wang, J. Chinese Patent: 201911249189.7, 2019.

44. Yao, Y.; Xin, J.; Li, Y.; Wei, D.; Wang, J. Chinese Patent: 201911248840.9, 2019.

45. Li, Y.; Xin, J.; Liu, H.; Wei, D.; Wang, J.; Yao, Y. Chinese Patent: 201910371013.2, 2019.

46. Li, Y.; Xin, J.; Du, X.; Wei, D.; Yao, Y.; Wang, J. Chinese Patent: 201910371432.6, 2019.

47. Yao, Y.; Xin, J.; Li, Y.; Wei, D.; Wang, J. Patent: PCT/CN2020/078140, 2020.

48. Mamone, S.; Rezaei-Ghaleh, N.; Opazo, F.; Griesinger, C.; Gloggler, S., Singlet-filtered NMR spectroscopy. Sci Adv 2020, 6 (8), eaaz1955.

49. Stevanato, G.; Hill-Cousins, J. T.; Håkansson, P.; Roy, S. S.; Brown, L. J.; Brown, R. C.; Pileio, G.; Levitt, M. H., A nuclear singlet lifetime of more than one hour in room-temperature solution. Angew. Chem. Int. Ed. 2015, 54 (12), 3740–3743.

50. Levitt, M. H., Long live the singlet state! J. Magn. Reson. 2019, 306, 69–74.

51. Tayler, M. C.; Levitt, M. H., Singlet nuclear magnetic resonance of nearly-equivalent spins. PCCP 2011, 13 (13), 5556–5560.

52. DeVience, S. J.; Walsworth, R. L.; Rosen, M. S., Preparation of nuclear spin singlet states using spin-lock induced crossing. PhRvL 2013, 111 (17), 173002.

53. Pileio, G.; Carravetta, M.; Hughes, E.; Levitt, M. H., The long-lived nuclear singlet state of 15N-nitrous oxide in solution. J. Am. Chem. Soc. 2008, 130 (38), 12582–12583.

54. Ahuja, P.; Sarkar, R.; Vasos, P. R.; Bodenhausen, G., Long-lived States in Multiple-Spin Systems. Chemphyschem 2009, 10 (13), 2217–2220.

55. Pileio, G.; Concistrè, M.; Carravetta, M.; Levitt, M. H., Long-lived nuclear spin states in the solution NMR of four-spin systems. J. Magn. Reson. 2006, 182 (2), 353–357.

56. Vinogradov, E.; Grant, A. K., Hyperpolarized long-lived states in solution NMR: Three-spin case study in low field. J. Magn. Reson. 2008, 194 (1), 46–57.

57. Grant, A. K.; Vinogradov, E., Long-lived states in solution NMR: Theoretical examples in three-and four-spin systems. J. Magn. Reson. 2008, 193 (2), 177–190.

58. Pileio, G.; Levitt, M. H., J-Stabilization of singlet states in the solution NMR of multiple-spin systems. J. Magn. Reson. 2007, 187 (1), 141–145.

59. Tayler, M. C.; Marie, S.; Ganesan, A.; Levitt, M. H., Determination of molecular torsion angles using nuclear singlet relaxation. J. Am. Chem. Soc. 2010, 132 (24), 8225–8227.

