## Supplementary text; Materials and methods; Figures S1 to S6; Tables S1 to S2 for "Molecularly Targeted Magnetic Resonance Imaging and Spectroscopy"

**This PDF file includes:**

Supplementary text
Materials and methods
Figures S1 to S6
Tables S1 to S2

**Supplementary Information text**

Spin evolution in YAWX-NMR

Figure S1 is the scheme of the pulse sequence, YAWX-NMR. The density matrix analysis of this pulse sequence is given in the following(1).

- **2-spin system (AGG):**

**The initial state** $\rho_0 = \hat{I}_{1z} + \hat{I}_{2z}$

⇓ $\frac{\pi}{2}(\hat{I}_{1y} + \hat{I}_{2y})$

① $\rho_1 = \hat{I}_{1x} + \hat{I}_{2x}$

⇓ $\hat{H}, \tau_{SL,max}$

② $\rho_2 = \frac{1}{2}\left(\hat{I}_{1x} + \hat{I}_{2x}\right) + 2\hat{I}_{1x}\hat{I}_{2x} + \hat{I}_{1y}\hat{I}_{2y} + \hat{I}_{1z}\hat{I}_{2z}$

⇓ $g_1$

③ $\rho_3 = 2\hat{I}_{1x}\hat{I}_{2x} + \hat{I}_{1y}\hat{I}_{2y} + \hat{I}_{1z}\hat{I}_{2z}$

⇓ Decoupling

④ $\rho_4 = S_0 + others$

⇓ $g_2$

⑤ $\rho_5 = S_0$

⇓ $\hat{H}, \tau_{SL,max}$

⑥ $\rho_6 = \frac{1}{2}(\hat{I}_{1x} + \hat{I}_{2x} + 2\hat{I}_{1x}\hat{I}_{2x} + 2\hat{I}_{1z}\hat{I}_{2z} + E)$

Where

$$\hat{H} = \omega_1 \hat{I}_{1z} + \omega_2 \hat{I}_{2z} + 2\pi J_{12} \hat{I}_1 \cdot \hat{I}_2 + 2\pi J_0 (\hat{I}_{1x} + \hat{I}_{2x})$$

$$J_0 = J_{12}$$

$$\tau_{SL,max} = \frac{1}{\Delta v \sqrt{2}} = \frac{0.707}{\Delta v}$$

$$S_0 = \frac{E}{2} - \hat{I}_{1x}\hat{I}_{2x} - \hat{I}_{1y}\hat{I}_{2y} - \hat{I}_{1z}\hat{I}_{2z}$$

*others*: the coherences which can be eliminated by gradient pulses.

- **3-spin system ($H^a$, $H^b$, $H^{b'}$ of NAA and $H^a$, $H^b$, $H^d$ of DA):**

**The initial state** $\rho_0 = \hat{I}_{1z} + \hat{I}_{2z} + \hat{I}_{3z}$

$\frac{\pi}{2}(\hat{I}_{1y} + \hat{I}_{2y} + \hat{I}_{3y})$ $\Downarrow$

① $\rho_1 = \hat{I}_{1x} + \hat{I}_{2x} + \hat{I}_{3x}$

$\hat{H}, \tau_{SL,max}$ $\Downarrow$

② $\rho_2 = \sqrt{3}\hat{I}_{1x} + \hat{I}_{1y} + \hat{I}_{2x} + \hat{I}_{3x} + 2\hat{I}_2 \cdot \hat{I}_3$

$g_1$ $\Downarrow$

③ $\rho_3 = 2\hat{I}_2 \cdot \hat{I}_3$

Decoupling $\Downarrow$

④ $\rho_4 = Q + others$

$g_2$ $\Downarrow$

⑤ $\rho_5 = Q$

$$\hat{H}, \tau_{SL,max} \Downarrow$$

⑥ $$\rho_6 = \hat{I}_{2x} + \hat{I}_{3x} + 2\hat{I}_{2x}\hat{I}_{3x}$$

Where

$$\hat{H} = \omega_1\hat{I}_{1z} + \omega_2\hat{I}_{2z} + \omega_3\hat{I}_{3z} + 2\pi(J_{12}\hat{I}_1 \cdot \hat{I}_2 + J_{13}\hat{I}_1 \cdot \hat{I}_3 + J_{23}\hat{I}_2 \cdot \hat{I}_3) + 2\pi J_0(\hat{I}_{2x} + \hat{I}_{3x})$$

$$J_0 = J_{23}$$

$$\tau_{SL,max} = \frac{1}{\Delta v\sqrt{2}} = \frac{0.707}{\Delta v}$$

$$\mathrm{Q} = \frac{2}{\sqrt{3}}(\hat{I}_1 \cdot \hat{I}_2 + \hat{I}_2 \cdot \hat{I}_3 + \hat{I}_1 \cdot \hat{I}_3)$$

*others*: the coherences which can be eliminated by gradient pulses.

Optimization of the spin lock in YAWX-NMR/MRI/MRS by using the Matlab program

The spin system in AGG: For AGG, the two-spin system consisting of $H^c$ and $H^{c'}$ is exploited to create the singlet state.

The Hamiltonian of a two-spin system, $\hat{H}_2$, is as follows:

$$\hat{H}_2 = \omega_1\hat{I}_{1z} + \omega_2\hat{I}_{2z} + 2\pi J_{12}\hat{I}_1 \cdot \hat{I}_2 + 2\pi J_0(\hat{I}_{1x} + \hat{I}_{2x})$$

By choosing a suitable radiation frequency, the resonance frequencies and J couplings of this spin system are: $\omega_1 = -2.85Hz,\ \omega_2 = 2.85Hz, J_{12} = -18.36Hz$. In the YAWX-NMR pulse, the x direction of spinlock pulse is the key to prepare the two-spin singlet state. Under the Hamiltonian, $\hat{H}_2$, the density matrix of the spin system will transfer from $\hat{I}_{1x} + \hat{I}_{2x}$ into multi-quantum states including a singlet term. The transfer efficiency from $\hat{I}_{1x} + \hat{I}_{2x}$ to the singlet depends on the spinlock time $\tau_{SL}$ and the power level, $J_0$. To optimize the singlet transfer efficiency, a Matlab program has been made to simulate the transfer process.

Figure S3 is the contour plot of the singlet transfer efficiency against the spinlock time $\tau_{SL}$ and the power level of spin lock.

We have made the similar simulation for the three-spin system(2) of NAA (i.e., $H^a$ , $H^b$ and $H^{b'}$). The resonance frequencies and J couplings used in the simulation are as follows: $\omega_1 = -895.5Hz, \omega_2 = -4Hz, \omega_3 = 4Hz, J_{12} = 5.6Hz, J_{13} = 6.2Hz, J_{23} = 17.2Hz$ . The contour plot of the singlet transfer efficiency against the spinlock time $\tau_{SL}$ and the power level of spin lock is shown in Figure S4.

### Materials and Methods

#### Materials

All reagents are used as received without further purification. N-acetylaspartate (NAA, 99%, Sigma-Aldrich), Dopamine hydrochloride (DA, 98%, Aladdin), a tripeptide (H-Ala-Gly-Gly-OH, AGG, 98.89%, Nanjing Peptide Biotech Ltd.), insulin (CAS: 11070-73-8, Sigma I-5500, Sigma), deuterium oxide (99.9 atom % D, CIL). The AGG, NAA, and DA aqueous solutions are prepared by dissolving the corresponding solutes into $D_2O$. The concentration of each solution sample is about 6 mg/ml.

#### Methods

NMR experiments

$^1$H NMR experiments were performed at a 500 MHz Bruker instrument. A Bruker TBI probe equipped with a three dimensional gradient was utilized in the experiments. The experimental temperature was 25 ℃. The recycle delay (TR) in the experiments was set to 10 s to ensure the full relaxation of the signals. The detailed parameters used in the NMR experiments are listed in Table S1. The power level and duration time of the spin lock were optimized before the experiment. A MATLAB program was designed for this purpose.

MRI and MRS experiments

MRI and MRS experiments were performed at a 500 MHz Bruker instrument. A Bruker TBI probe equipped with a three dimensional gradient was utilized in the experiments. The $^{1}H$ spin echo MRI image of the sample in Figure 3 and 4 was acquired by using a standard $^{1}H$ spin echo imaging sequence ($T_E$ = 15.5 ms, $T_R$ = 10 s, field of view = 5.6 × 5.6 $mm^2$). The molecularly targeted images of the sample in Figure 3 was acquired by using the pulse sequence shown in Figure 1b. In these experiments, the power level and duration time of the spin lock were optimized before the MRI/MRS experiment. A Matlab program was designed for this purpose. This program is provided on request. A set of detailed parameters used in the MRI/MRS experiments are listed in Table S2. The recycle delay (TR) in the experiments was set to 10 s to ensure the full relaxation of the signals. The experimental temperature was 25 ℃.

The sample used for the MRI/MRS experiments were prepared as follows: Firstly, four capillary tubes (1 mm diameter, ~0.1 mm thickness), containing water (60 % $D_2O$ and 40% $H_2O$) and the NAA, AGG, and DA aqueous solution, were prepared. Then the four capillary tubes were carefully loaded into the 5 mm glass tube containing water (40% $D_2O$ and 60% $H_2O$).

**Fig. S1:** The scheme of the pulse sequence, YAWX-NMR. The different steps are marked by the different numbers.

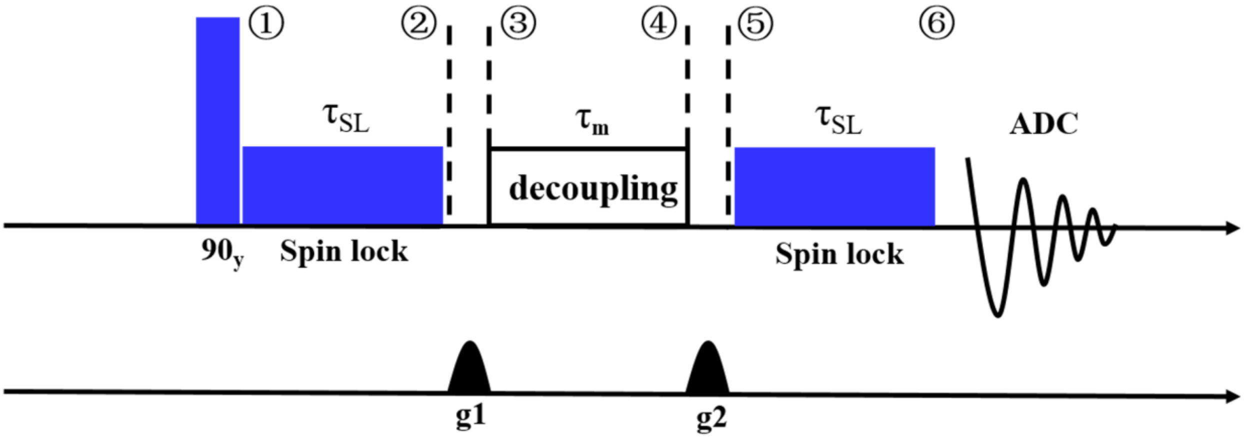

**Fig. S2:** The $^{1}H$ NMR spectra of the aqueous solution of a tripeptide (L-Alanine-glycine-glycine, AGG), acquired by using: a) the single pulse excitation sequence; b) the YAWX-NMR pulse sequence to select the signals of $H^{c}$ and $H^{c'}$ of AGG. The $^{1}H$ NMR spectra of the aqueous solution of N-acetylaspartate (NAA), acquired by using: c) the single pulse excitation sequence; d) the YAWX-NMR pulse sequence to select the signals of $H^{b}$ and $H^{b'}$ of NAA. The $^{1}H$ NMR spectra of the aqueous solution of dopamine (DA), acquired by using: e) the single pulse excitation sequence; f) the YAWX-NMR pulse sequence to select the signals of $H^{d}$ of DA. The experimental temperature is room temperature. The recycle delay in the experiments was set to 10 s to ensure the full relaxation of the signals.

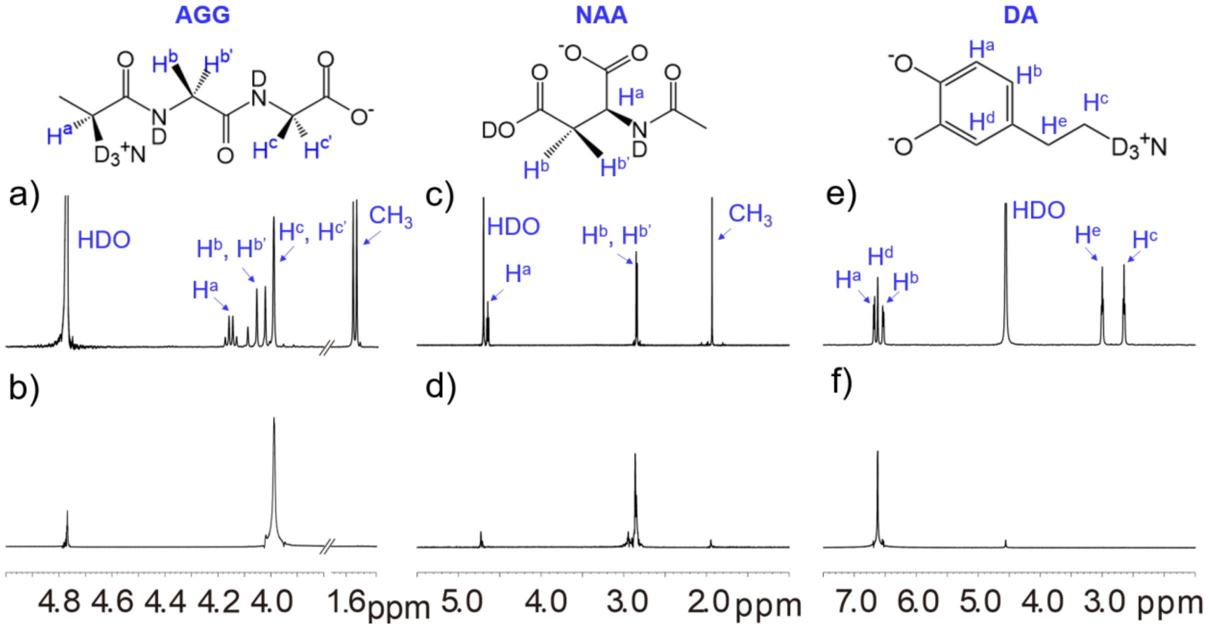

**Fig. S3:** The contour plot of the singlet transfer efficiency against the spinlock time $\tau_{SL}$ and the power level of spin lock. The simulation parameters are from the two-spin system of AGG.

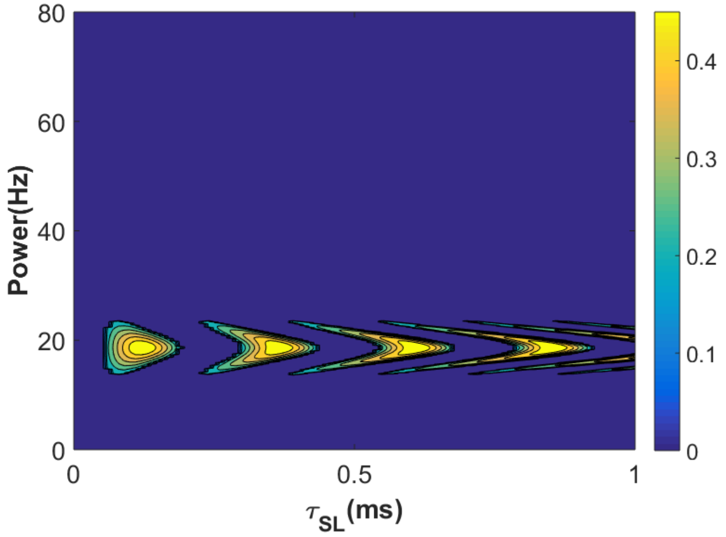

**Fig. S4:** The contour plot of the singlet transfer efficiency against the spinlock time $\boldsymbol{\tau_{SL}}$ and the power level of spin lock. The simulation parameters are from the three-spin system of NAA ($H^a$ , $H^b$ and $H^{b'}$).

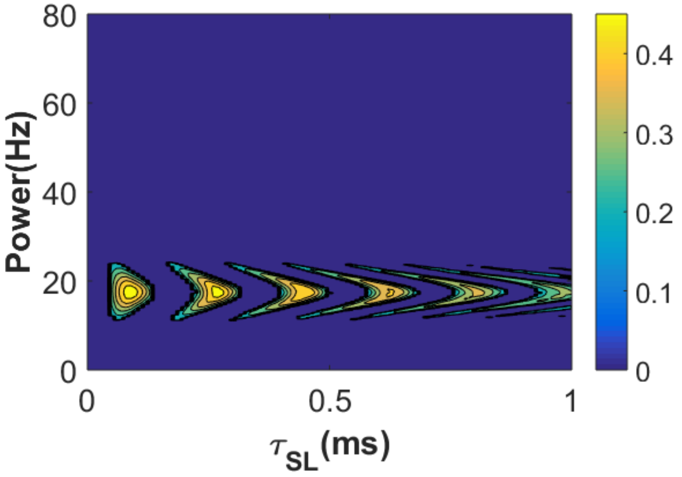

**Fig. S5:** The contour plot of the conversion efficiency of singlet state of AGG ($H^c$ and $H^{c'}$) against the chemical shift difference between $H^c$ and $H^{c'}$ ($\Delta\omega$) and J coupling.

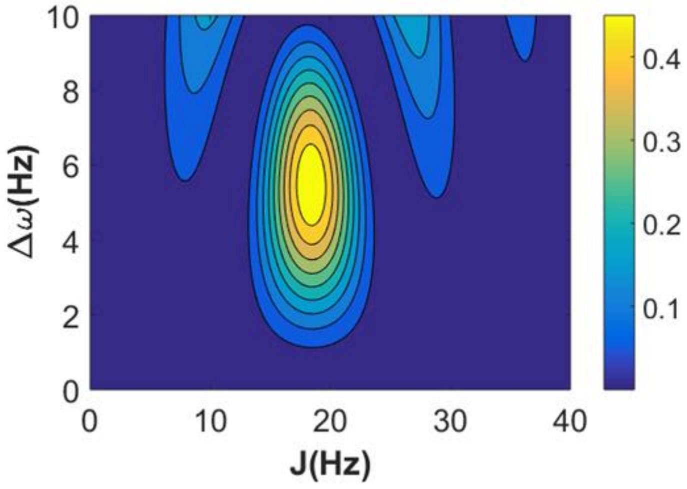

**Fig. S6:** a) The $^1$H NMR spectra acquired by using a) the single pulse excitation pulse sequence (UP) and the YAWX-NMR pulse sequence (DOWN). The sample was the DA aqueous solution with the concentration of 26.1 mM. These two spectra were obtained after a single scan. b) The $^1$H NMR spectra acquired by using a) the single pulse excitation pulse sequence (UP) and the YAWX-NMR pulse sequence (DOWN). The sample was the DA aqueous solution with the concentration of 26.1 μM(3). (DOWN). These two spectra were obtained after accumulation of 512 scans. $H^d$ of DA molecule were selectively probed in the experiments. All of the experiments were performed at room temperature.

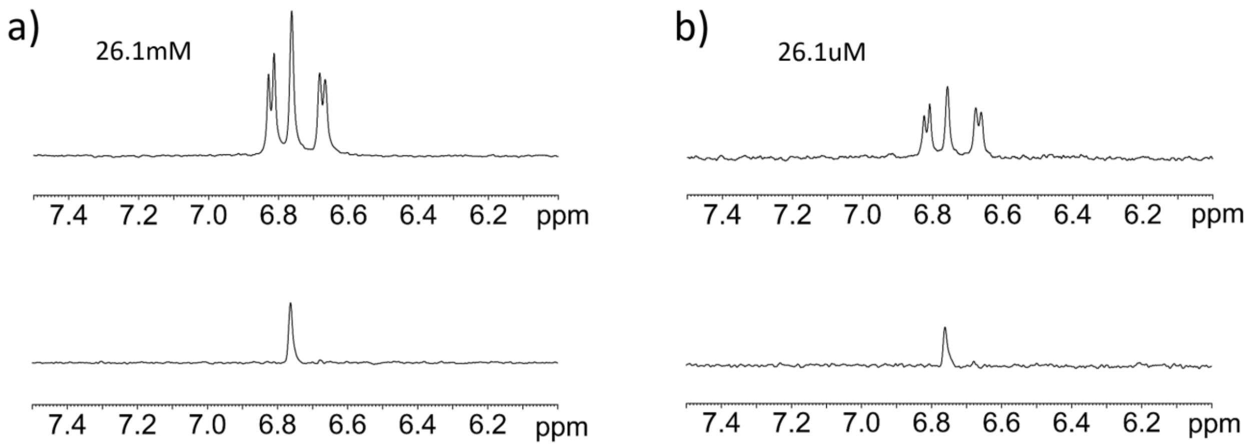

**Table S1:** The parameters used in the YAWX-NMR experiments

| | AGG | NAA | DA |
|---|---|---|---|
| **The first 90$^o$ pulse time (μs)** | **7.75** | **7.75** | **7.75** |
| **Spin lock duration (ms)** | **40** | **40** | **27** |
| **Spin lock power (Hz)** | **6.86** | **6.86** | **4.87** |
| **Decoupling time (ms)** | **18** | **18** | **18** |
| **Decoupling power (Hz)** | **105** | **66** | **92** |
| **Z gradient (G/cm)** | **27** | **27** | **27** |

**Table S2:** The parameters used in the YAWX-MRI/MRS experiments

| | AGG | NAA | DA |
|---|---|---|---|
| **The first 90$^o$ pulse time (μs)** | **7.75** | **7.75** | **7.75** |
| **Spin lock duration (ms)** | **70** | **70** | **40** |
| **Spin lock power (Hz)** | **18.5** | **18.5** | **10** |
| **Decoupling time (ms)** | **18** | **18** | **18** |
| **Decoupling power (Hz)** | **105** | **66** | **92** |
| **Sinc wave time (ms)** | **1** | **1** | **1** |
| **Sinc wave power (Hz)** | **500** | **500** | **500** |
| **x gradient (G/cm)** | **27** | **27** | **27** |
| **y gradient (G/cm)** | **15** | **15** | **15** |
| **z gradient (G/cm)** | **0.527** | **0.527** | **0.527** |

### SI References


1. S. J. DeVience, R. L. Walsworth, M. S. Rosen, Preparation of nuclear spin singlet states using spin-lock induced crossing. *Phys Rev Lett* **111**, 173002 (2013).
2. P. Ahuja, R. Sarkar, P. R. Vasos, G. Bodenhausen, Long-lived states in multiple-spin systems. *Chemphyschem* **10**, 2217-2220 (2009).
3. T. Lee, L. X. Cai, V. S. Lelyveld, A. Hai, A. Jasanoff, Molecular-level functional magnetic resonance imaging of dopaminergic signaling. *Science* **344**, 533-535 (2014).
